# HIV-1 uncoating occurs via a series of rapid biomechanical changes in the core related to individual stages of reverse transcription

**DOI:** 10.1101/2021.01.16.426924

**Authors:** Sanela Rankovic, Akshay Deshpande, Shimon Harel, Christopher Aiken, Itay Rousso

## Abstract

The HIV core consists of the viral genome and associated proteins encased by a cone-shaped protein shell termed the capsid. Successful infection requires reverse transcription of the viral genome and disassembly of the capsid shell within a cell in a process known as uncoating. The integrity of the viral capsid is critical for reverse transcription, yet the viral capsid must be breached to release the nascent viral DNA prior to integration. We employed atomic force microscopy to study the stiffness changes in HIV-1 cores during reverse transcription in vitro in reactions containing the capsid-stabilizing host metabolite IP_6_. Cores exhibited a series of stiffness spikes, with up to three spikes typically occurring between 10-30, 40-80, and 120-160 minutes after initiation of reverse transcription. Addition of the reverse transcriptase (RT) inhibitor efavirenz eliminated the appearance of these spikes and the subsequent disassembly of the capsid, thus establishing that both result from reverse transcription. Using timed addition of efavirenz, and analysis of an RNAseH-defective RT mutant, we established that the first stiffness spike requires minus-strand strong stop DNA synthesis, with subsequent spikes requiring later stages of reverse transcription. Additional rapid AFM imaging experiments revealed repeated morphological changes in cores that were temporally correlated with the observed stiffness spikes. Our study reveals discrete mechanical changes in the viral core that are likely related to specific stages of reverse transcription. Our results suggest that reverse-transcription-induced changes in the capsid progressively remodel the viral core to prime it for temporally accurate uncoating in target cells.

## Introduction

The HIV-1 core consists of single-stranded RNA (ssRNA) encapsulated within a cone-shaped protein shell called a capsid. Successful infection requires reverse transcription of the ssRNA into double-stranded DNA (dsDNA) - a complex process that requires the polymerase and RNaseH enzymatic activities of reverse transcriptase and is punctuated by two strand transfer events (1). The double-stranded viral DNA is then transported into the nucleus and is integrated into the target cell genome. As the intact HIV-1 core is too large to cross the nuclear pore (reviewed in (2)), the capsid is thought to disassemble prior to and/or during nuclear import (3, 4) in a process known as uncoating. Data from several studies suggest a complex relationship between HIV-1 capsid stability, reverse transcription, and uncoating (5–11). Specifically, while having a viral capsid of sufficient stability is important for reverse transcription, reverse transcription appears to promote uncoating in target cells. However, the detailed mechanism by which reverse transcription promotes HIV-1 uncoating, and the detailed physical changes that the core undergoes during reverse transcription, are essentially undefined.

Several studies have suggested that HIV-1 uncoating is regulated by cellular factors and metabolites. Inositol hexakisphosphate (IP_6_) is an abundant cellular metabolite that is present in all mammalian cells (12). IP_6_ is incorporated into HIV-1 virions during assembly at a level of approximately 300 molecules per virion (13), which appears to be important for particle assembly and for stabilization of the mature viral capsid. IP_6_ binds to the positively charged pores of hexamers of the HIV-1 capsid protein (CA) and promotes assembly of CA into conical structures *in vitro* (13–15). Addition of IP_6_ to permeabilized HIV-1 particles also stabilizes the viral capsid, resulting in protection of viral DNA from degradation by added nuclease *in vitro* (16–18).

Atomic force microscopy (AFM) provides a unique approach to monitor the physical changes in the HIV-1 core that occur during reverse transcription. AFM is nondestructive and can be applied to unfrozen, unfixed samples, thus permitting many consecutive measurements on single cores over extended time periods. AFM uses a physical probe to obtain measurements of particle stiffness, a mechanical property, and can also be operated in an imaging mode to obtain information about particle shape and structure. Recently, we used AFM to observe the physical changes in the HIV-1 core occurring during uncoating *in vitro*. We found that addition of capsid-stabilizing ligands, including the host protein cyclophilin A, the small molecule antiviral compound PF74, and the cell metabolite IP_6_, increased the stiffness of cores (18–20). Analysis of the stiffness and morphology of isolated cores undergoing reverse transcription revealed that viral core stiffness reaches a maximum 5–7 hours into the reaction, subsequently resulting in a breach in the narrow end of the capsid and its ultimate dissociation (20, 21). Subsequently, we observed that treatment of HIV-1 cores with IP_6_ appeared to accelerate the process, resulting in more rapid uncoating and possibly also reverse transcription (18). Under these conditions, the previously observed single stiffness peak was not apparent, perhaps owing to the low temporal resolution of the averaged stiffness measurements (~2 h) employed in that study, which may have obscured the more rapid aspects of the reactions.

Recent studies have demonstrated that addition of IP_6_ promotes efficient reverse transcription within HIV-1 cores *in vitro* (22, 23). Concentrations of IP_6_ from 10 to 100 μM resulted in ~1000-fold increase in full-length minus strand synthesis, resulting in a high percentage of cores undergoing the second strand transfer step. These studies further linked the enhancement of reverse transcription to the capsid-stabilizing effects of IP_6_. In the present study, we employed AFM to define the biomechanical and morphological changes of individual isolated HIV-1 cores occurring during reverse transcription in the presence of IP_6_ *in vitro*. We observed a pattern of three rapid core stiffness spikes, which were dependent on reverse transcription. Pharmacological inhibition of reverse transcription at specific time points demonstrated that the second and third spikes require later stages of reverse transcription, as does the uncoating effect. Overall, our results show that reverse transcription is accompanied by a rapid, multi-step, mechanical stimulation process that is essential for complete core disassembly.

## Results

To study the effect of IP_6_ on reverse transcription-related morphological and mechanical changes in the HIV-1 core, we prepared samples for AFM analysis using a previously described method (20, 21). To obtain information about rapid changes in the core, we performed more frequent measurements on individual cores during reverse transcription. Acquiring the stiffness profile of an individual core during several hours of reverse transcription was technically demanding, because repetitive scanning of the same core by the AFM probe often causes the immobilized core to detach from the glass substrate. Despite this challenge, we acquired the stiffness profiles of 50 individual cores as they underwent reverse transcription under various conditions. In the first set of experiments, the stiffness profiles of 22 individual IP_6_-treated cores were acquired, of which 14 exhibited one or more stiffness spikes (Fig. 1 A-E and S1) and 8 showed no spikes (Fig. S2). Spikes that were larger than the background noise level were selected using a MatLab script that calculated their appearance time and magnitude (Fig. S3). Most IP_6_-treated cores exhibited multiple spikes, with 3 observed in most cases (9 out of 14, Fig. 1 F). Analysis of the spikes revealed some variability in their appearance times, yet they occurred at three distinct time frames (Fig. S1-O). Notably, the first spike appeared between the initial 10-30 minutes and was typically followed by a second and third spike, which appeared after 40-80 and 120-160 minutes, respectively. In one core, we observed a long spike, which is probably two stiffness spikes that were not resolved, and thus a total of two spikes were determined for this trajectory (Fig. 1 D). Of the 14 active cores analyzed, only one core had a profile consisting of a single clear spike (Fig. 1 E). The magnitudes of the spikes were variable, possibly owing to intrinsic variation in capsid shape and/or stability. However, their normalized magnitudes (peak value divided by the spike baseline value) are quite reproducible with an average value of 2.2±0.1 (n=36, Fig, S1 P).

**Fig. 1.**
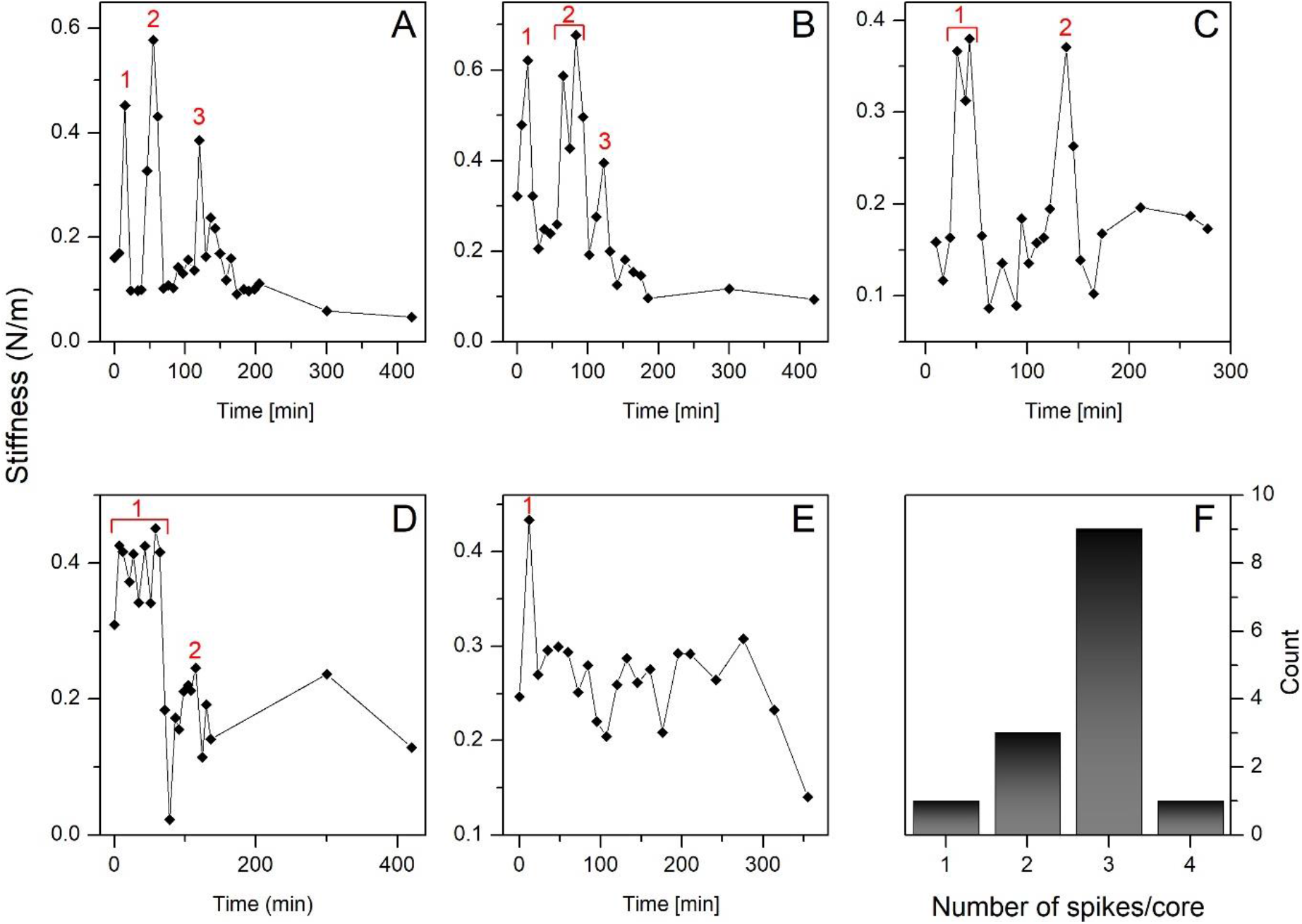
Stiffness trajectories of five individual isolated HIV-1 cores treated with 100 μM IP_6_ as a function of the progress of reverse transcription. Cores were adhered to HMDS-coated glass slides and kept in MOPS buffer. Reverse transcription was initiated by the addition of dNTPs and MgCl_2_ to the cores, and their stiffness values were measured using AFM. (A-E) Stiffness profiles of individual HIV-1 cores undergoing reverse transcription. Stiffness spikes are labeled by a red number. (F) The distribution of number of stiffness spikes per individual core.

With AFM operating in the quantitative imaging (QI) mode, we next analyzed the morphology of the cores during the multi-stiffness spikes pattern (i.e. in the first ~3 hours of reverse transcription. We observed clear changes in the shape of the cores that coincided temporally with the observed stiffness spikes (Fig. 2 and supporting video). To quantify these changes, we calculated a parameter known as the circularity index of the core perimeter. A circularity value of one indicates a perfect circle, whereas lower values describe elongated objects. Interestingly, we observed that the changes in the circularity index of the core occurred at similar times as the stiffness spikes, suggesting that both occurred in parallel (Fig. 2 A-D). In agreement with our quantified shape analysis, topographic AFM images reveal that the core acquired a more circular shape during each mechanical excitation and then rapidly returned to its initial morphology (Fig. 2 E and supporting videos).

**Fig. 2.**
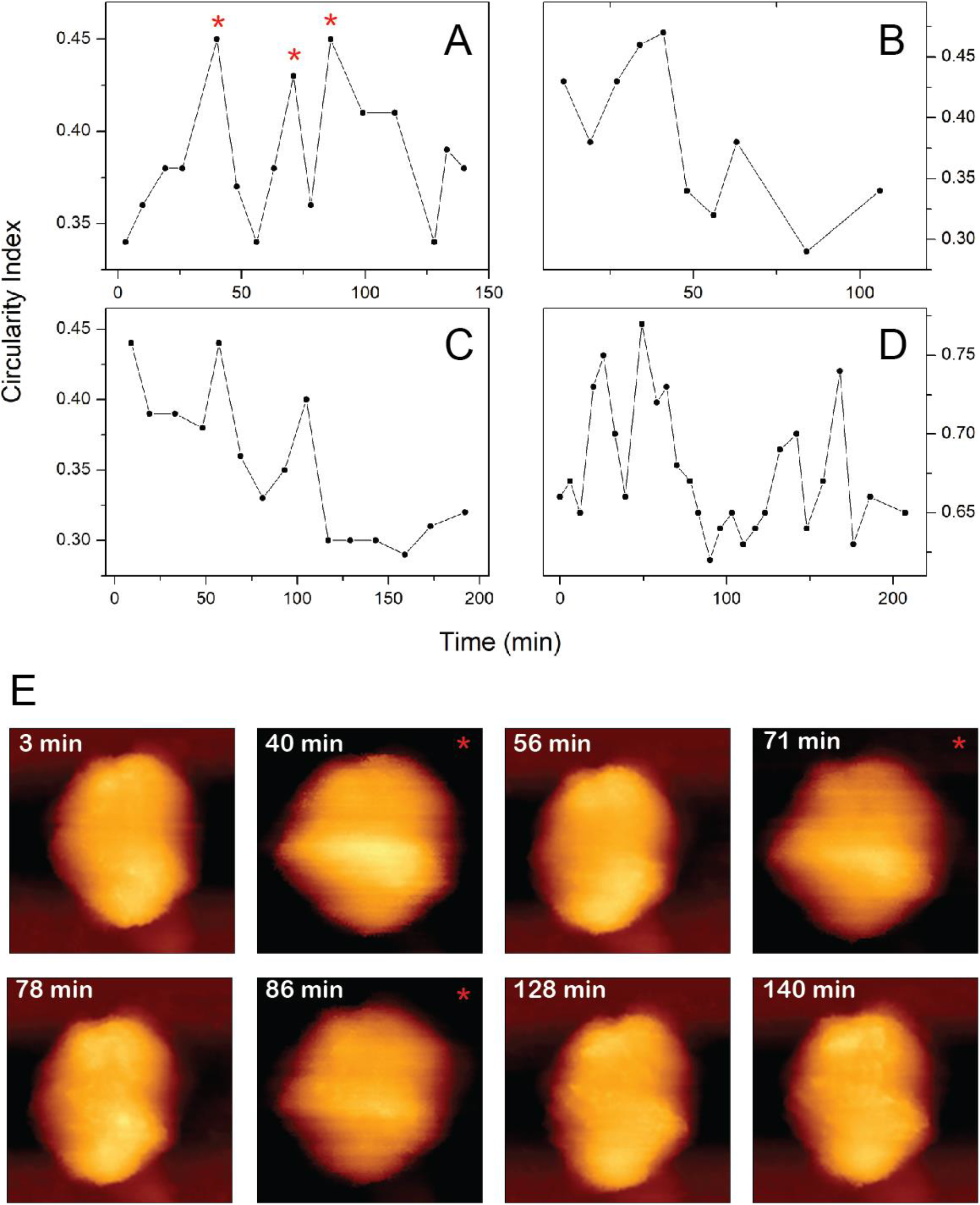
The morphological changes that HIV-1 cores treated with 100 μM IP_6_ undergo during the first 3 hours of reverse transcription. **(A-D)** Circularity index of four HIV-1 cores imaged by AFM (out of a total of six) plotted against reverse transcription time. Cores were adhered to HMDS-coated glass slides and kept under MOPS buffer. For reference, the circularity indexes of equilateral triangle, square and hexagon, are 0.6, 0.78 and 0.9, respectively. (E) Representative time frames taken from the time-lapse movie showing the morphology of the core (panel A) during reverse transcription. The images of the core at the corresponding circularity index peaks are labeled with a red star.

To determine the relationship between the stiffness spikes and DNA synthesis, we inhibited reverse transcription by adding efavirenz to the reactions. When added at the beginning of the reaction, no spikes were observed and the stiffness of the core remained relatively constant during the reaction (Fig. 3 A and S4). Moreover, the number of intact cores per μm^2^ of grid area remained unchanged during the experiment (a total of 21 cores were imaged). Based on these observations, we conclude that occurrence of core stiffness spikes and capsid disassembly both depend on DNA synthesis.

**Fig. 3.**
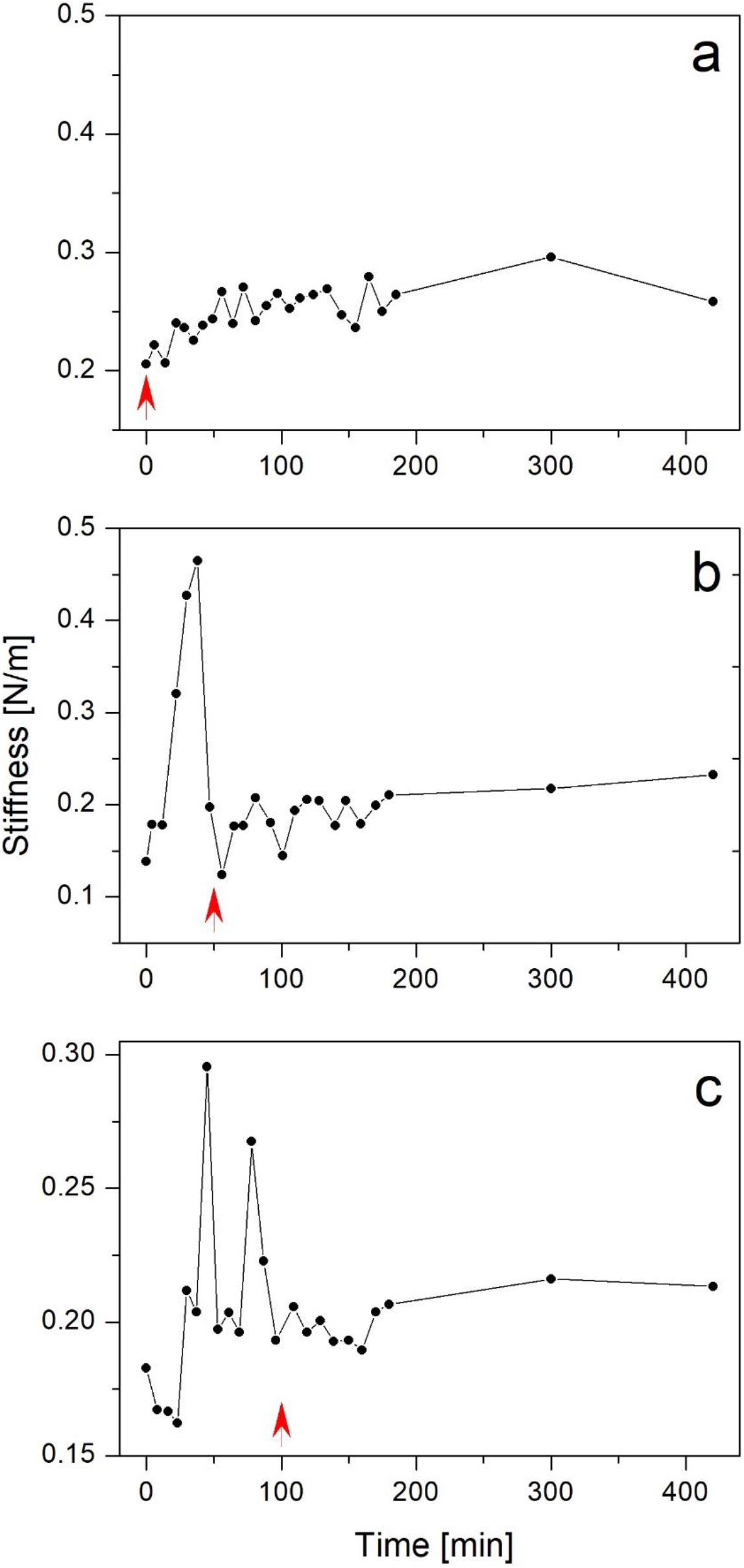
Effect of inhibiting reverse transcription on the stiffness trajectories of individual isolated HIV-1 cores treated with 100 μM IP_6_ as a function of the progress of reverse transcription. Stiffness of individual HIV-1 cores quantified every 8 min by AFM during reverse transcription. The initial average stiffness value of cores prior to reverse transcription is represented as time zero. (A-C) Addition of efavirenz (red arrow) to cores after 0, 50 and 100 minutes of reverse transcription, respectively.

In additional experiments, we added efavirenz at specific time points during reverse transcription to determine the effects on occurrence of the second and third spikes. Each experiment ran for 7 hours. Addition of efavirenz after 50 minutes resulted in a single stiffness peak, which corresponded temporally to the earliest peak observed in reactions lacking the inhibitor (Fig. 3B and S5). A comparison of the numbers of observed intact cores per given area at the beginning and end of the experiment showed that nearly all cores remained intact (a total of 23 cores were imaged), indicated that uncoating was also inhibited.

Addition of efavirenz to the reactions after 100 min resulted in two stiffness spikes (Fig. 3 C and S6, a total of 6 individual cores were analyzed from which 2 core did not exhibit spikes) at times corresponding to the first and second spikes in uninhibited reactions. Strikingly, morphological analysis of the cores, obtained by AFM at the end of the experiment (after 7 hours), showed that nearly all of the 23 cores imaged underwent partial disassembly but did not progress to complete core disassembly (Fig. 4). Consistently with our previous findings (20, 21), apparent openings in the capsid (the size of which varied between cores) were localized at or near the narrow end of the core (in 20 of 23 cores imaged after reverse transcription).

**Fig. 4.**
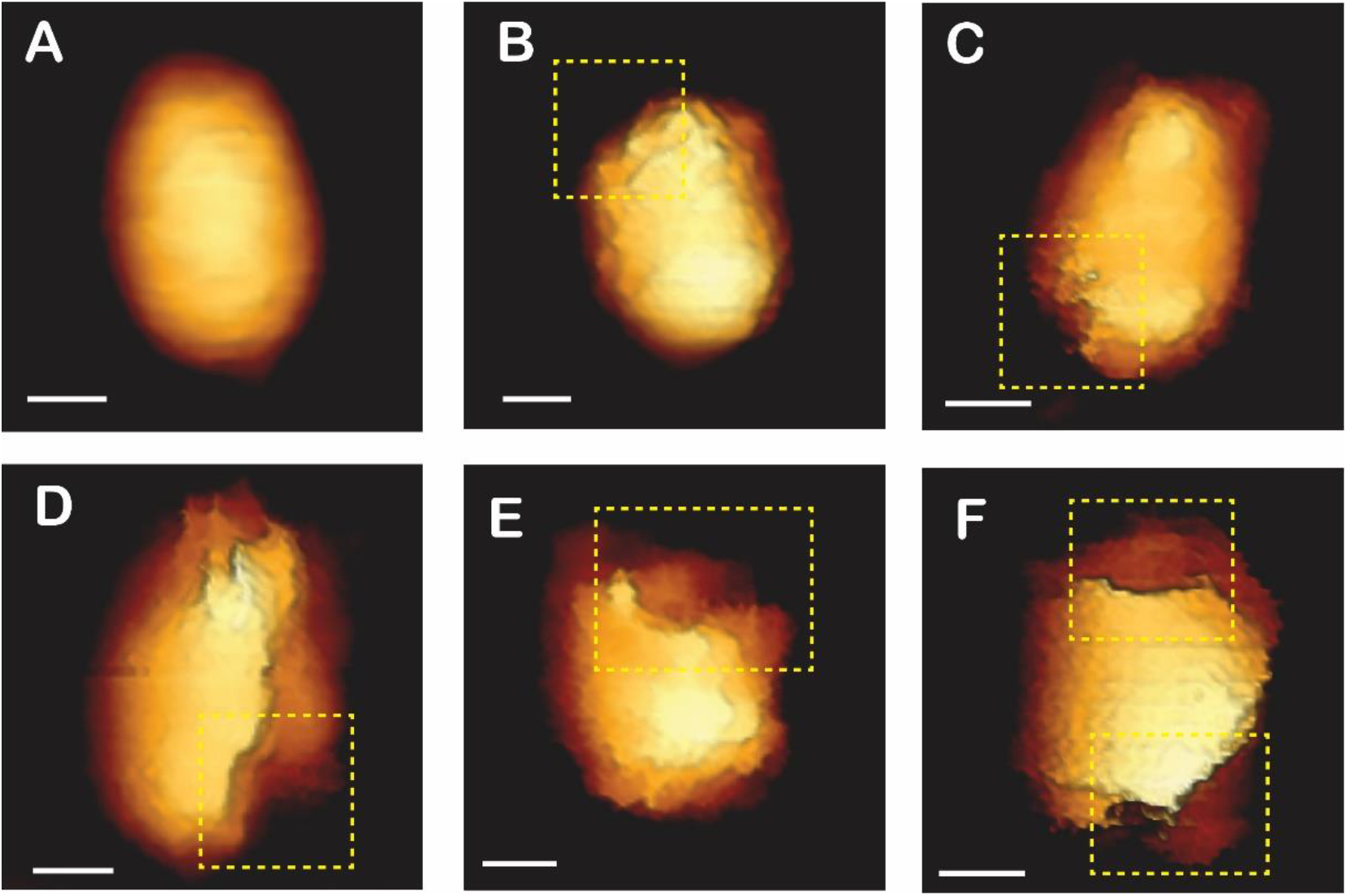
Effect of inhibiting reverse transcription on the morphological changes that HIV-1 cores treated with 100 μM IP_6_ undergo during reverse transcription. Cores were imaged by AFM during reverse transcription, which was inhibited by addition of efavirenz after 100 minutes. Topographic images were acquired using AFM operated in quantitative imaging mode. (A) A representative native HIV-1 core observed prior to reverse transcription (out of a total of 18 cores that were imaged). (B-F) Deformed and damaged core visualized after 7 hours of reverse transcription. For clarity, openings in the cores are shown within a dashed yellow rectangle. Scale bars, 50 nm. A total of 23 cores were visualized during reverse transcription.

We also tested the effects of addition of efavirenz at time points after all three stiffness spikes had occurred. Core morphology was measured twice subsequently to efavirenz addition, near and at the end of the experiment (after 5 and 7 hours, respectively). Similar to adding efavirenz after 100 minutes, its addition following 130 minutes resulted in nearly all cores (22 of 29 cores visualized) undergoing partial rather than complete disassembly (4 cores remained intact and thus likely did not undergo reverse transcription). Inhibiting reverse transcription after 4 hours produced a mixture of partially (~60%) and completely (~30%) disassembled cores (of the total of 14 cores imaged). Finally, we terminated reverse transcription after 5 hours. Morphological analysis of the samples 2 hours later revealed that nearly all cores had disassembled completely.

The appearance of multiple spikes in core stiffness suggested the existence of multiple intermediate steps and/or stalling phases (24) during reverse transcription. To further understand the cause of these stiffness spikes, we examined cores isolated from the RNAseH-defective RT mutant E478Q. In this mutant, reverse transcription cannot proceed beyond the synthesis of the first 180 nucleotides from the 5’ end of genomic RNA (this step is commonly referred to as minus-strand strong stop synthesis) owing to a failure to degrade the RNA template and anneal to the 3’ end of the genome (1). Intriguingly, analysis of individual E478Q cores during reverse transcription revealed that their mechanical trajectories exhibited only a single prominent spike, which appeared ~20 minutes into reverse transcription (Fig. 5 A and S7, a total of 8 individual cores were analyzed from which 2 did not exhibited stiffness spikes). Moreover, the cores remained intact throughout the entire duration of the experiment (a total of 20 cores were imaged). Thus, our AFM results indicate that the multiple spikes in the core stiffness trajectories correspond to discrete steps in the reverse transcription process, with the first peak linked to minus-strand strong stop synthesis.

**Fig. 5.**
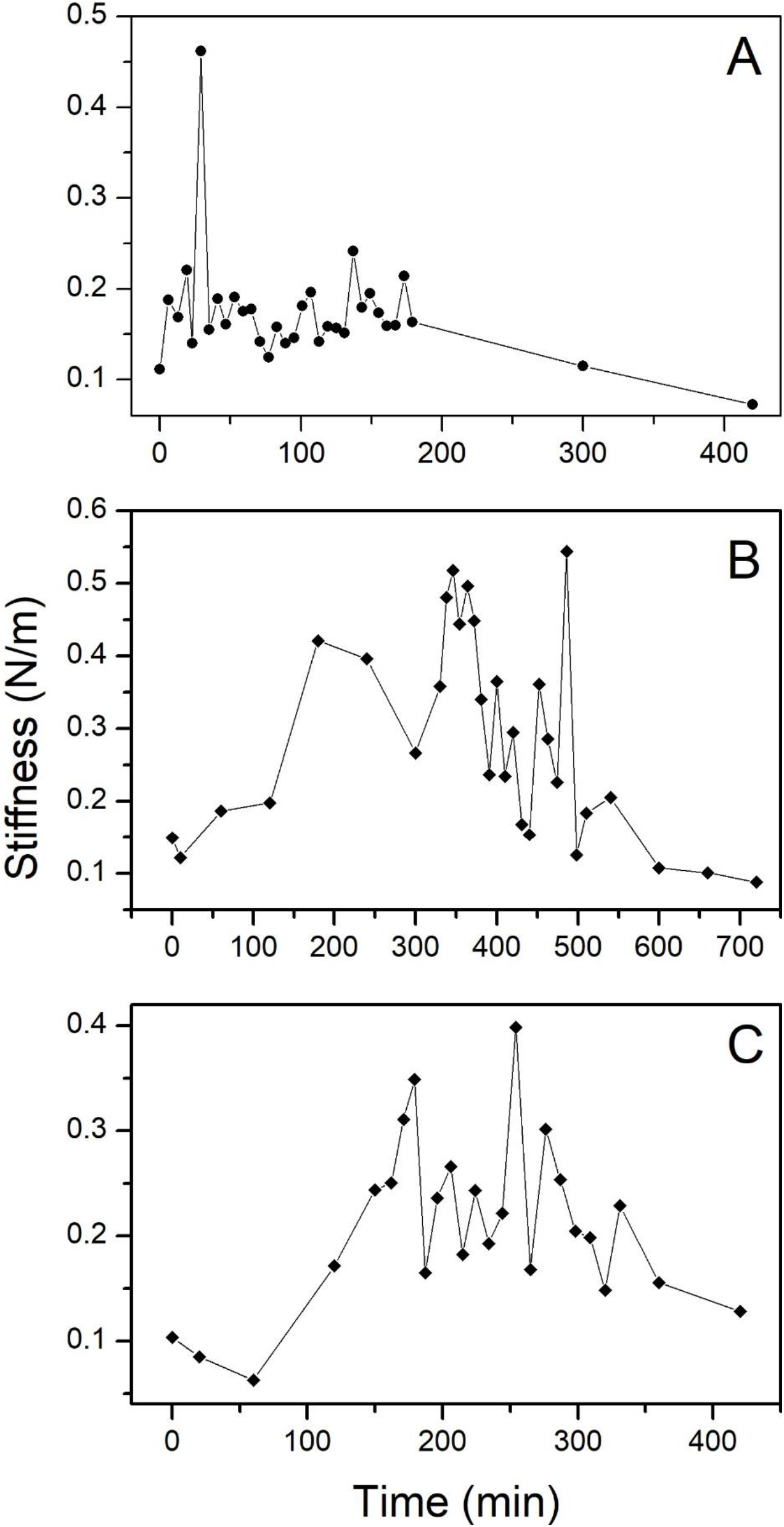
Stiffness trajectories of individual RNAse H-defective and non IP_6_ treated HIV-1 cores during reverse transcription. Cores were adhered to HMDS-coated glass slides and kept in MOPS buffer. Each panel shows the profile of a single core. (A) Stiffness of RNAse H-defective HIV-1 cores in reaction containing IP_6_. Stiffness trajectory of individual WT (B) and E45A (C) HIV-1 cores during reverse transcription in the absence of added IP_6_.

Lastly, we analyzed cores isolated from wild type and hyperstable capsid mutant (E45A) viruses during reverse transcription in the absence of added IP_6_ to determine whether the multiple peak stiffness trajectories are an IP_6_-induced effect or represent a normal feature of reverse transcription. Under our experimental conditions, the stiffness trajectories induced by reverse transcription were significantly slower in untreated cores compared with IP_6_-treated cores (19–24 hours compared with 7 hours, respectively). At these lengthy time scales, it is a daunting task to analyze individual cores with high temporal resolution. We therefore focused on a 3-hour window centered around the observed stiffness peak (which occurred after 4 and 7 hours of transcription for E45A and WT, respectively), with an individual core analyzed every ~8 minutes within this time course. Remarkably, the mechanical trajectories of individual E45A and WT cores exhibited multiple stiffness spikes during reverse transcription, which appeared after 2.5–7.5 hours of reverse transcription of WT and after 2.5–4.5 hours for E45A (Fig. 5 B-C and S8, a total of 5 WT and 6 E45A cores were analyzed from which 2 and 3, respectively, did not exhibited stiffness spikes). Taken together, these results indicate that the multiple spikes in the stiffness trajectories is a common feature in HIV-1 reverse transcription.

## Discussion

The stability of the viral capsid is critical for HIV-1 reverse transcription and productive uncoating. Here we employed AFM to reveal biomechanical aspects of HIV-1 uncoating by evaluating the effect of the capsid-stabilizing cellular metabolite IP_6_ on the mechanical stability and morphology of isolated HIV-1 cores during reverse transcription. By analyzing individual cores every 6–8 minutes during the course of reverse transcription, we observed that the majority of the cores examined exhibited three stiffness spikes. In one of the 14 cores, we detected a single stiffness spike during the reaction. Intriguingly, the reverse transcription stiffness profiles shared similar features. Specifically, the spikes occurred at three time windows during the course of reverse transcription. The first stiffness spike appeared during the first 10-20 minutes of reverse transcription, and the second and third spikes appeared during 4-80 and 120-160 minutes, respectively. When a reverse transcriptase inhibitor was added to the buffer, the stiffness of the cores did not change and the cores remained intact throughout the measurement (up to 12 h). In addition, out of the 22 IP_6_-treated cores analyzed, only 14 (64%) exhibited stiffness spikes and underwent disassembly. This is in agreement with the reported reverse transcription efficiency for IP_6_-treated cores (22, 23) and suggests that most but not all of the cores are active in reverse transcription. We therefore conclude that the observed three-spike profile that accompanies capsid disassembly depends on reverse transcription.

The observed biomechanical effects of reverse transcription observed in this study are likely to be physiologically relevant. IP_6_ is present in substantial concentrations in mammalian cells, and it is also enriched in HIV-1 particles via binding to Gag during virus assembly. However, it was the addition of exogenous IP_6_ to the reverse transcription reactions that initially revealed the series of stiffness spikes during reverse transcription, raising the possibility of a biochemical artifact. Subsequent analysis of the stiffness of HIV-1 cores from wild type and E45A mutant virions during reverse transcription in the absence of added IP_6_ clearly revealed a pattern of multiple stiffness spikes occurring a time frame within which we observed a single peak in our previous studies. We therefore conclude that reverse transcription itself induces a series of mechanical stimulations in native HIV-1 cores that are required for their disassembly *in vitro*.

In previous studies, we also observed mechanical and morphological changes in purified HIV-1 cores during the course of reverse transcription (20, 21). We found that core stiffness increased during reverse transcription, reaching a maximum after 7 hours, followed by complete disassembly of the capsid after 19–24 hours of transcription. Surprisingly, IP_6_-treated cores underwent complete disassembly after only 7 hours (18). We proposed that the effect of IP_6_ is mediated via binding to the positively charged central pore in CA hexamers in the capsid (13). In the absence of IP_6_, transport of the negatively charged dNTPs across the capsid shell through the positively charged pores may be attenuated owing to electrostatic interactions with the central hexamer channel. Binding between the six negative charges on IP_6_ and the six positive charges on the CA hexamer reduces the overall net positive charge of the pore, potentially facilitating influx of dNTPs and accelerating reverse transcription. This may explain the observed acceleration in reverse transcription-induced core disassembly. Surprisingly, averaged measurements of IP_6_-treated cores did not exhibit a stiffness peak during reverse transcription. We now think this is a result of the relatively poor temporal resolution of the measurement (~2 hr), which did not allow detection of faster changes, i.e. those observed in the present study.

The unique characteristics of the observed stiffness profiles raise an intriguing question: what does each individual stiffness spike represent? We suggest that each results from a specific stage of DNA synthesis, with the first peak occurring during synthesis of minus strand strong stop DNA. This is supported by our observation that cores from the RNaseH-defective mutant exhibited the first spike but not the subsequent ones. The first spike did not require release of the viral DNA from the RNA genome nor its annealing to the 3’ end of the genome, both of which depend on RNAseH activity. Therefore, capsid stiffening appears to be driven by DNA synthesis. This could result from the structural changes in the interior of the core that likely occur during reverse transcription, which are yet to be defined. It could also result from the process of DNA synthesis itself, which result in a long polyanionic molecule which could interact with the inner wall of the viral capsid, possibly stiffening it. Such a reinforcement effect has been observed in the minute virus of mice (MVM), where interactions between the viral DNA and the capsid increased the stiffness of the core (25). The view that the stiffness spikes correspond to specific stages of DNA synthesis is also supported by a report showing that the initial minus strand synthesis and transfer requires approximately 5 minutes in vitro (26). In that study, complete synthesis of the minus strand required approximately 105 ± 30 minutes, whereas the plus strand synthesis and transfer required an additional ~30 minutes. Thus, there appears to be temporal correspondence between stages DNA synthesis and occurrence of the observed core stiffness spikes. An additional possibility is that pyrophosphate molecules, which are a by-product of DNA synthesis resulting from dNTP hydrolysis, could interact with the capsid and alter its mechanical properties. Pyrophosphate could then dissociate, resulting in a return of the stiffness to baseline. Alternatively, interactions of pyrophosphate with the capsid could reduce the stiffness to baseline by competing with the nascent DNA for binding to the capsid wall, resulting in dissociation of the latter. In any case, it is intriguing that the core-stiffening effect of reverse transcription is apparent reversible, as reflected in the rapid decline of the stiffness following each spike.

Our observations (summarized in Fig. 6) reveal new insights into the mechanism of reverse transcription-induced uncoating. We observed a gap of 3–5 hours between mechanical perturbations (completed after two hours of reverse transcription) and the complete disassembly of the core (after 5–7 hours of transcription). In a previous study, we also detected a time gap in reactions lacking IP_6_, which was longer (stiffness spike occurring at 7 hours and complete disassembly after 19–24 hours of reverse transcription) (21). During the stiffness spikes, AFM imaging analysis suggested that the morphology of the core remained intact. However, the core appears to change its shape during the biomechanical phase of reverse transcription. We propose a “cumulative damage” model in which individual steps in reverse transcription result in subtle damage to the capsid lattice, priming the core for later uncoating required for integration. When the core undergoes a single mechanical stimulation, the subsequent damage appears to be too minor to propagate into detectable structural damage. By contrast, two spikes results in damage that results in partial capsid disassembly. Similarly, when the core experiences three mechanical stimulations, but reverse transcription is then inhibited, the core also undergoes only partial disassembly. The structural damage results in complete capsid disassembly only when all three stiffness spikes occur and reverse transcription proceeds beyond the period of mechanical stimulation. These results suggest that reverse transcription can be divided into two phases: the first phase is characterized by the induction of a series of mechanical stimulations whereas the second phase is mechanically silent but critical for complete core disassembly. When reverse transcription is terminated after 4 hours, the accumulated damage produces a mixture of partially and completely disassembled cores. By contrast, blocking reverse transcription after 5 hours does not affect core disassembly, such that fully disassembled cores are obtained at the end of the experiment (7 hours). Based on our results, we propose that multi-step mechanical excitation induces cumulative structural damage that is required for reverse transcription-induced core disassembly. Complete core disassembly requires 5 hours of reverse transcription, within which the last 3 hours are mechanically silent. The accumulated damaged following 5 hours of reverse transcription is large enough to proceed into full core disassembly that is completed during the final 2 hours and occurs independently of continued reverse transcription.

**Fig. 6.**
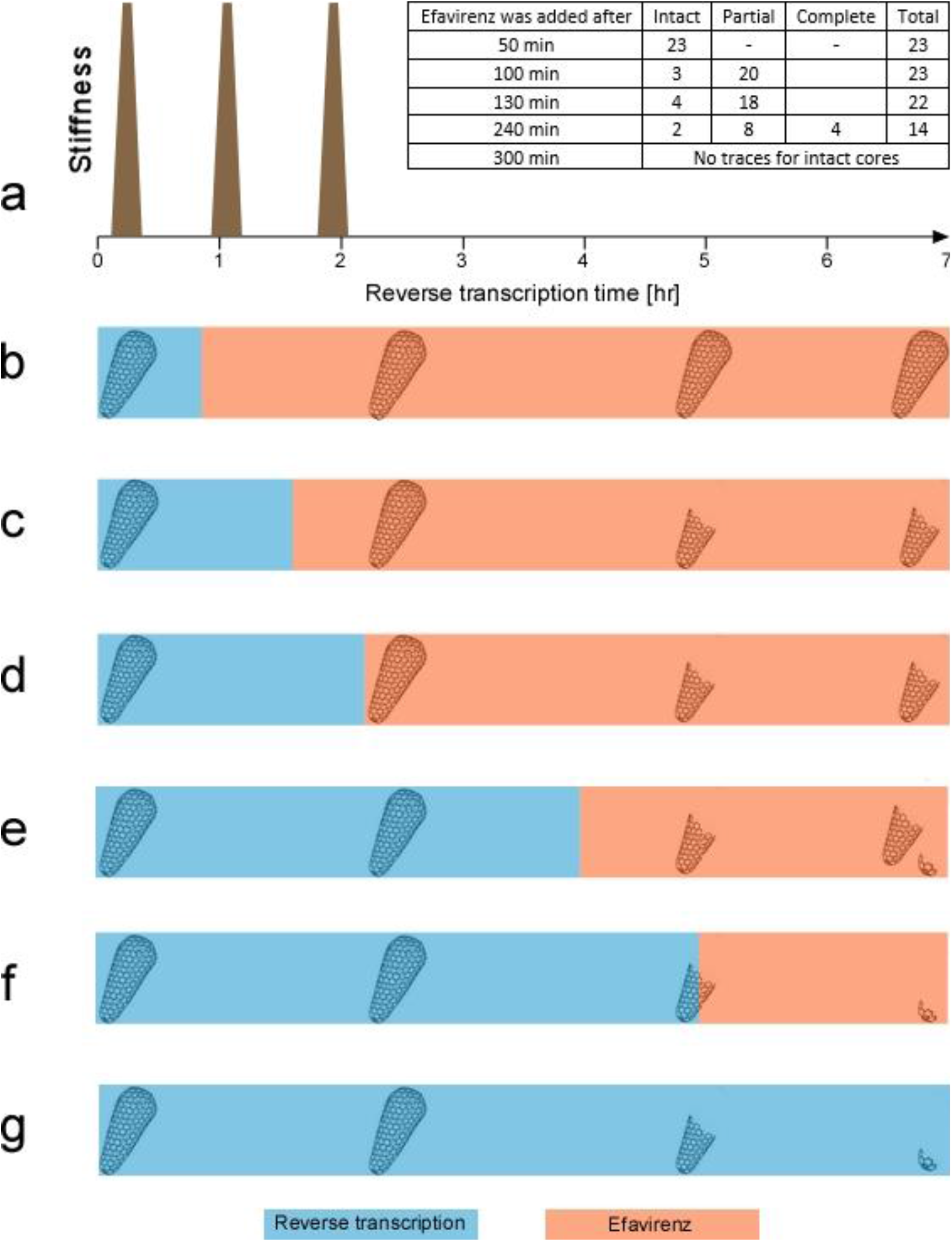
A schematic representation of the effect of efavirenz addition at specific time points during reverse transcription on the morphology of the core. The morphology of the core is presented as intact, partially disassembled, or completely disassembled. (a) A cartoon representing the trajectory of the mechanical events occurring during reverse transcription (RT), including the three observed stiffness spikes (brown). (b–g) The morphology of the core during 7-hour experiments in which the RT inhibitor efavirenz was added after (b) 50 min (c) 100 min, (d) 130 min, (e) 4 h, (f) 5 h, and (g) 7 h of reverse transcription. The RT period is shown in blue whereas the period during which RT was inhibited by efavirenz is shown in orange.

Recent work suggests that HIV-1 reverse transcription and complete uncoating occur inside the nucleus (27, 28). Moreover, these processes are highly likely to be regulated by host factors which highlight the potential for perturbation by capsid-binding small molecule antivirals such as PF74 (29). Our “cumulative damage” model is also compatible with uncoating in the nucleus, with early (i.e. cytoplasmic) reverse transcription-induced mechanical stresses to the capsid prime the core for later uncoating and integration, which may be triggered by binding of host factors in the nuclear pore complex and/or within the nucleus.

## Materials and Methods

### Isolation of HIV-1 cores

HIV-1 pseudovirions were used for the isolation of HIV-1 cores. Pseudovirus particles were produced by a previously described protocol (20, 21, 30). Briefly, approximately 10^6^ human embryonic kidney (HEK) 293T cells were transfected with 2.5 μg of ΔEnv IN- HIV-1 plasmid (DHIV3-GFP-D116G) (31) using 10 μg of polyethylenimine (PEI, branched, MW ~25,000, Sigma-Aldrich). After 20 h, the medium was replaced with fresh medium (Dulbecco modified Eagle medium supplemented with 10% heat-inactivated bovine serum, 1% penicillin-streptomycin, and 1% glutamine), and the cells were incubated at 37°C in 5% CO2. After 6 h, the supernatant was harvested, centrifuged at 1,000 rpm for 10 min, and filtered through a 0.45-μm-pore size filter. The virus-containing supernatant was concentrated by ultracentrifugation in a SW-28 rotor (25,000 rpm, 2 h, 4°C) using OptiPrep density gradient medium (Sigma-Aldrich). Part of the supernatant containing pelleted viruses was collected, mixed with 10 ml of TNE buffer (50 mM Tris-HCl, 100 mM NaCl, 0.1 mM EDTA [pH 7.4]) and added to the 100-kDa molecular mass cutoff Vivaspin 20 centrifugal concentrators (100,000 MWCO, Sartorius AG, Germany). The mixture was centrifuged twice at 2,500 × *g* for 25 to 30 min at 4°C, until the supernatant level in the concentrators reached 200–300 μl.

From concentrated virus-containing supernatant, viral cores were isolated using a previously described protocol (32) with modifications. Briefly, ∼40 μl of purified HIV-1 pseudovirus particles was mixed with an equal amount of 1% Triton-X diluted in 100 mM 3-(N-morpholino)propanesulfonic acid (MOPS) buffer (pH 7.0) and incubated for 2 min at 4 °C. The mixture was centrifuged at 13,800 × *g* for 8 min at 4°C. After removing the supernatant, the pellet was washed twice by adding ∼80 μl of MOPS buffer and centrifuging at 13,800 × *g* for 8 min at 4°C. The pallet was resuspended in 10 μl of MOPS buffer and used immediately for AFM studies.

### AFM measurements and analysis

AFM measurements and analysis were performed as previously described (20, 21). Briefly, 10 μl of isolated HIV-1 cores resuspended in MOPS buffer solution was incubated for 30 min at room temperature on hexamethyldisilazane (HMDS)-coated microscope glass slides (Sigma). Measurements were carried out in MOPS buffer, without sample fixation. Every experiment was repeated at least three times, each time with independently purified HIV-1 cores. IP_6_ was provided by the James laboratory as a generous gift. Measurements were carried out with a JPK Nanowizard Ultra-Speed atomic force microscope (JPK Instruments, Berlin, Germany) mounted on an inverted optical microscope (Axio Observer; Carl Zeiss, Heidelberg, Germany). Silicon nitride probes (mean cantilever spring constant, k_cant_=0.12 N/m, DNP, Bruker) were used. Height, topographic, and mechanical map images were acquired in quantitative imaging (QI) mode, at a rate of 0.5 lines/s and a loading force of 300 pN.

Core stiffness was obtained by the nanoindentation method, as previously described (20, 21, 30). Briefly, the stiffness value of each capsid was determined by acquiring ~400 force-distance (F-D) curves. To determine the stiffness value of cores, 20 F-D curves were obtained at rate of 20 Hz at each of 24 different points on the core surface. To confirm that the capsid or the core remained stable during the entire indentation experiment, we analyzed each experiment by plotting the individual measured point stiffness as a histogram, and as a function of the measurement count. Samples whose point stiffness values decreased consistently during experimentation were discarded, since this indicated that they had undergone irreversible deformation.

The maximum indentation of the sample was 4 nm, which corresponds to a maximum loading force of 0.2–1.5 nN. Prior to analysis, each curve was shifted to set the deflection in the noncontact section to zero. The set of F-D curves was then averaged. From the slope of the averaged F-D curve, measured stiffness was derived mathematically. The stiffness of the capsid was computed using Hooke’s law on the assumption that the experimental system may be modeled as two springs (the core and the cantilever) arranged in series. The spring constant of the cantilever was determined during experimentation by measuring thermal fluctuation (33). To reduce the error in the calculated point stiffness, we chose cantilevers such that the measured point stiffness was <70% of the cantilever spring constant.

### Endogenous reverse transcription (ERT) with WT and RT mutant viruses for AFM analysis

Reverse transcription was induced in cores attached to HMDS-coated microscope glass slides. To initiate reverse transcription, MOPS buffer was replaced with reverse transcription buffer (100 μM dNTPs and 1 mM MgCl_2_ in 100 mM MOPS buffer, pH=7.0) (34). To study the effect of the reverse transcription inhibitor (efavirenz, AIDS Reagent Program, NIH), efavirenz was added to the reverse transcription buffer to achieve a final concentration of 100 μM dNTPs, 1 mM MgCl_2_, and 100 nM. All measurements were carried out at room-temperature (23-25°C). IP_6_ was added to the reactions to a final concentration of 100 μM. Histograms of the individual measured point stiffness values derived from the consecutive force-distance curves of a single core at 7 reverse transcription time-points is shows in Fig. S9.

To analyze the single-core stiffness traces for spikes, the background for each trajectory was computed by first fitting a linear curve. Next, the averaged local minimum values was used to generate the background region by adding and subtracting this value to the linear curve (Fig. S3). Spikes are selected if their peak values are higher than the background. If a spike was split by a single measurement that was less than half of its height (as in Fig. 1 B and C), the two spikes were considered as one with an appearance time that was calculated as the averaged time of the two peaks.

Circularity index was calculated by dividing the area of the core multiplies by four pie by its squared perimeter. So that a perfect circle will give a circularity index of one and the lower the value is the, the less circular the shape of the core is. The circularity index of a perfect cone shaped core is 0.29. All data analysis was carried out using MATLAB software (The Math Works, Natick, MA).

## Acknowledgments

This work was supported in part by NIH grant R56 AI076121 and by the Israel Science Foundation (Grant 234/17). We thank Dr. Bryan Shepherd for helpful discussions. The following reagent was obtained from the NIH AIDS Reagent Program, Division of AIDS, NIAID, NIH: efavirenz.

