## Supplemental figures for "HIV-1 uncoating occurs via a series of rapid biomechanical changes in the core related to individual stages of reverse transcription"

### Slide 1
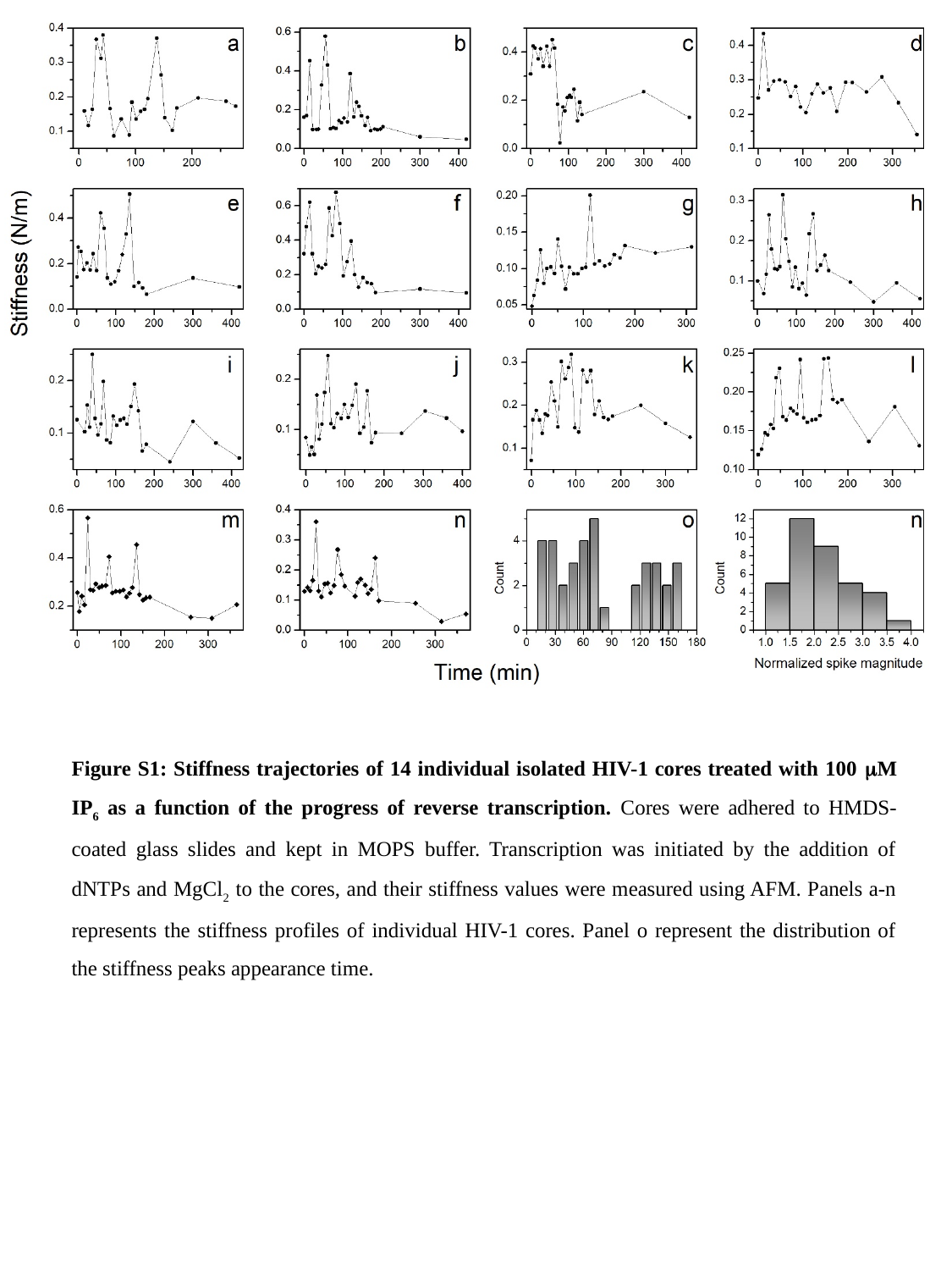

Figure S1: Stiffness trajectories of 14 individual isolated HIV-1 cores treated with 100 μM IP_6_ as a function of the progress of reverse transcription. Cores were adhered to HMDS-coated glass slides and kept in MOPS buffer. Transcription was initiated by the addition of dNTPs and MgCl_2_ to the cores, and their stiffness values were measured using AFM. Panels a-n represents the stiffness profiles of individual HIV-1 cores. Panel o represent the distribution of the stiffness peaks appearance time.

### Slide 2
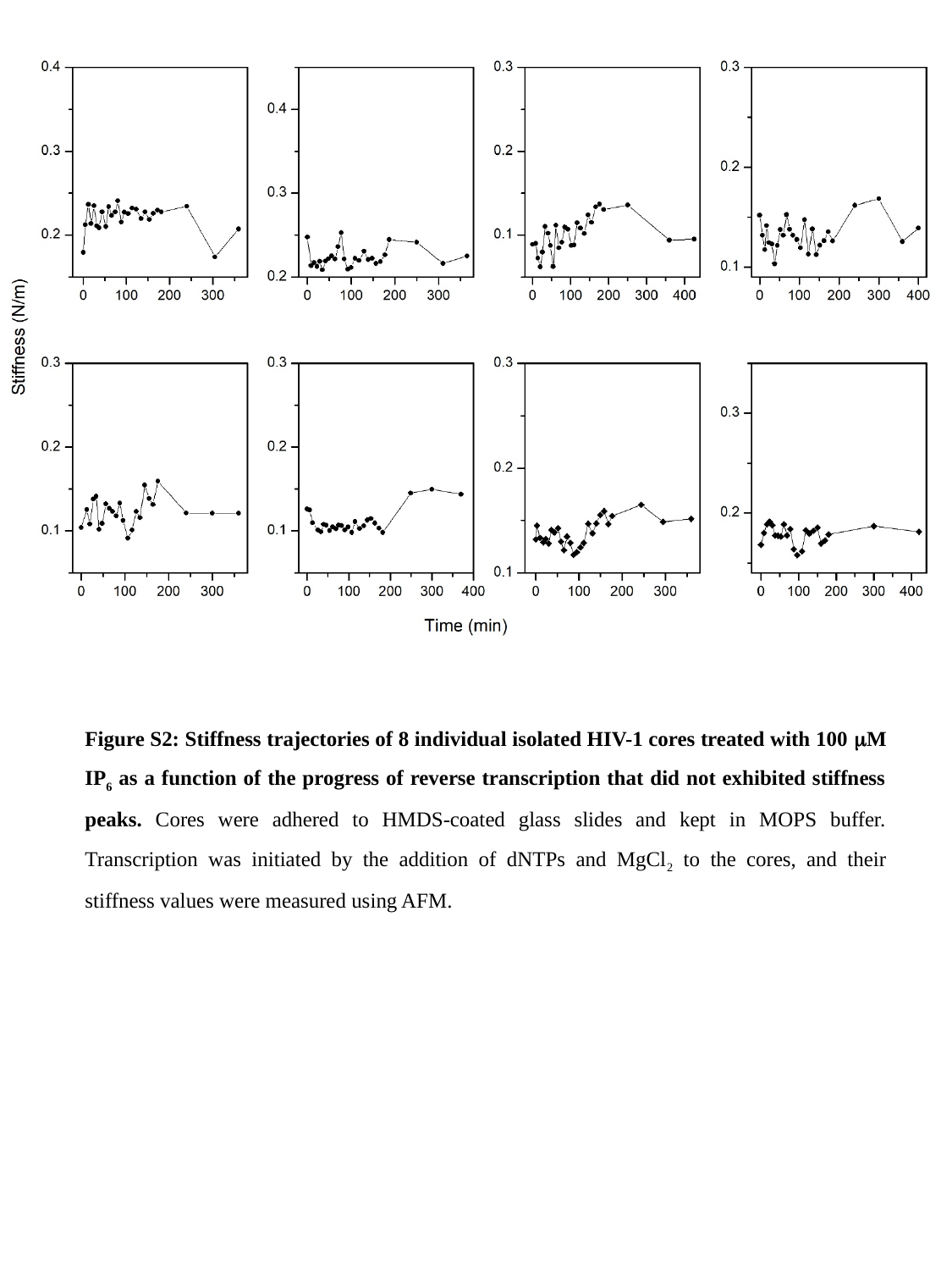

Figure S2: Stiffness trajectories of 8 individual isolated HIV-1 cores treated with 100 μM IP_6_ as a function of the progress of reverse transcription that did not exhibited stiffness peaks. Cores were adhered to HMDS-coated glass slides and kept in MOPS buffer. Transcription was initiated by the addition of dNTPs and MgCl_2_ to the cores, and their stiffness values were measured using AFM.

### Slide 3
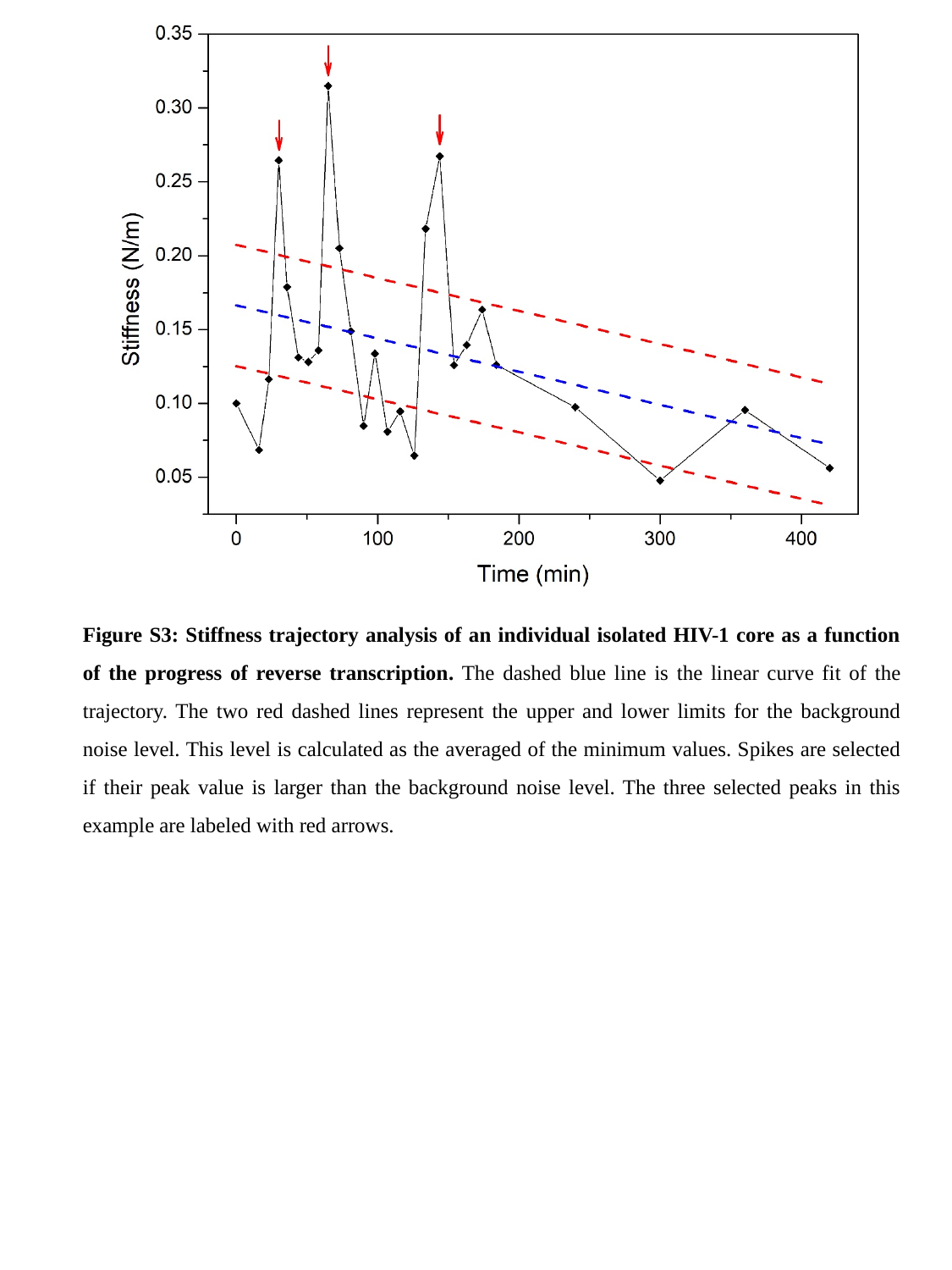

Figure S3: Stiffness trajectory analysis of an individual isolated HIV-1 core as a function of the progress of reverse transcription. The dashed blue line is the linear curve fit of the trajectory. The two red dashed lines represent the upper and lower limits for the background noise level. This level is calculated as the averaged of the minimum values. Spikes are selected if their peak value is larger than the background noise level. The three selected peaks in this example are labeled with red arrows.

### Slide 4
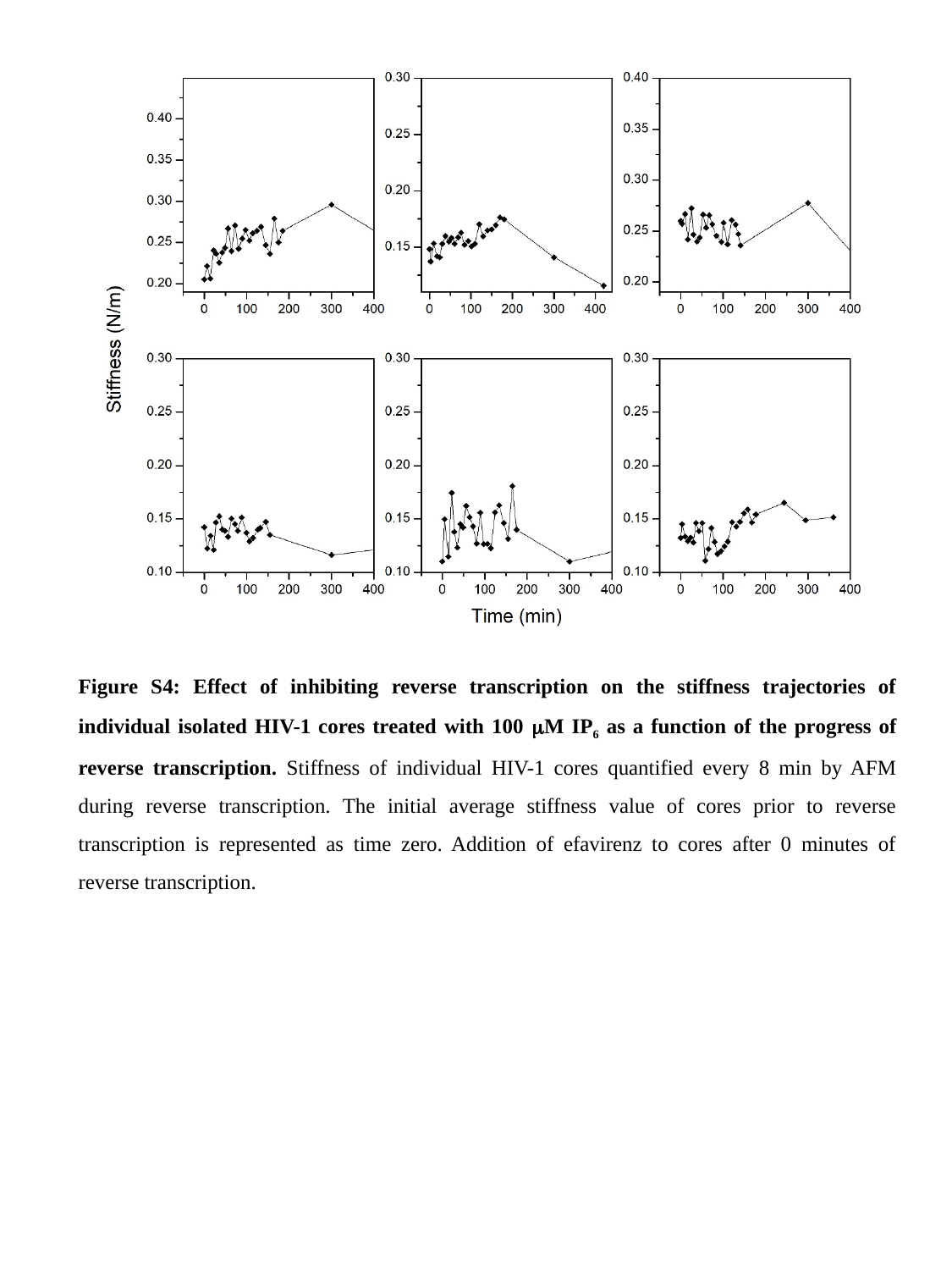

Figure S4: Effect of inhibiting reverse transcription on the stiffness trajectories of individual isolated HIV-1 cores treated with 100 μM IP_6_ as a function of the progress of reverse transcription. Stiffness of individual HIV-1 cores quantified every 8 min by AFM during reverse transcription. The initial average stiffness value of cores prior to reverse transcription is represented as time zero. Addition of efavirenz to cores after 0 minutes of reverse transcription.

### Slide 5
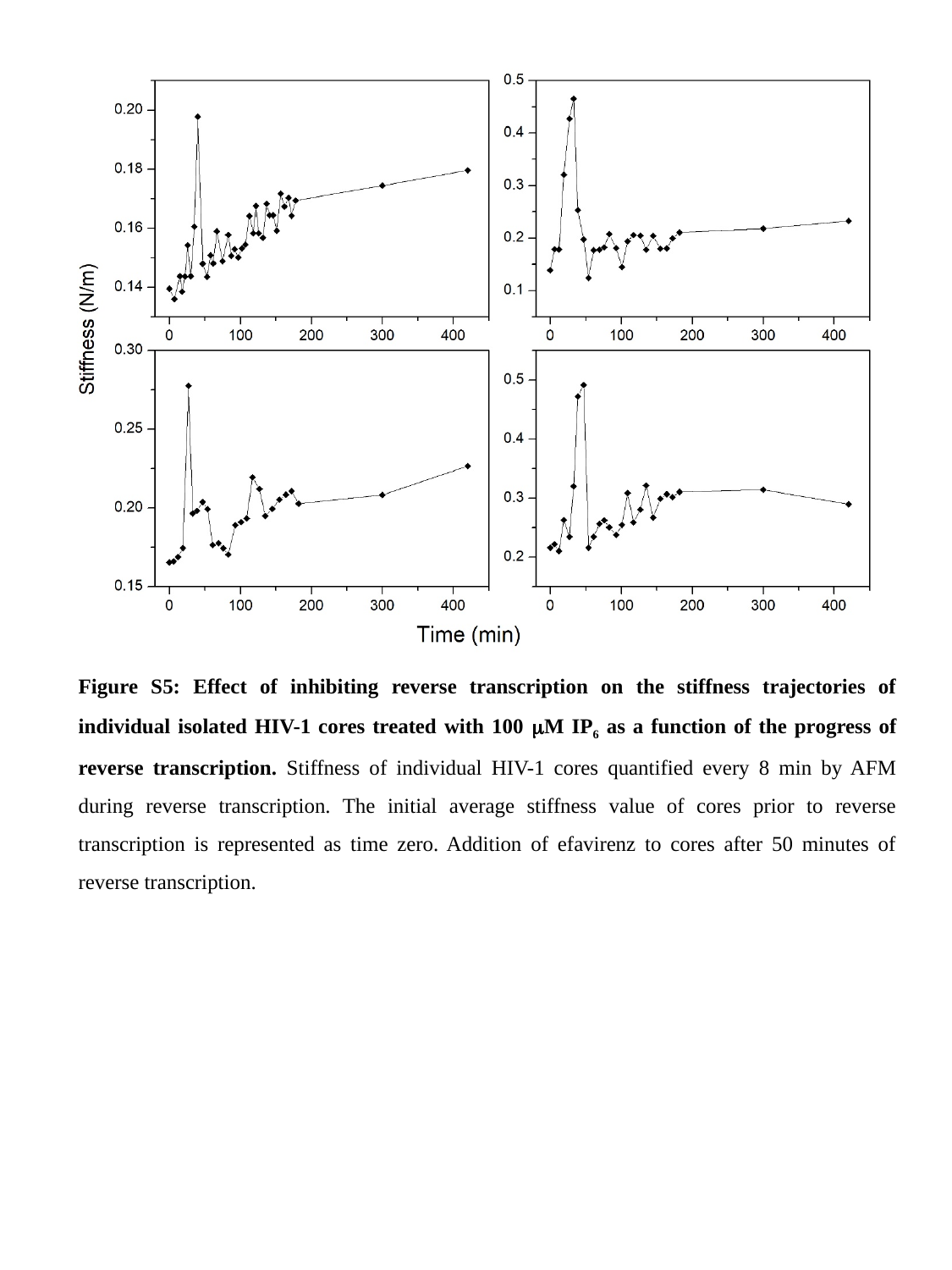

Figure S5: Effect of inhibiting reverse transcription on the stiffness trajectories of individual isolated HIV-1 cores treated with 100 μM IP_6_ as a function of the progress of reverse transcription. Stiffness of individual HIV-1 cores quantified every 8 min by AFM during reverse transcription. The initial average stiffness value of cores prior to reverse transcription is represented as time zero. Addition of efavirenz to cores after 50 minutes of reverse transcription.

### Slide 6
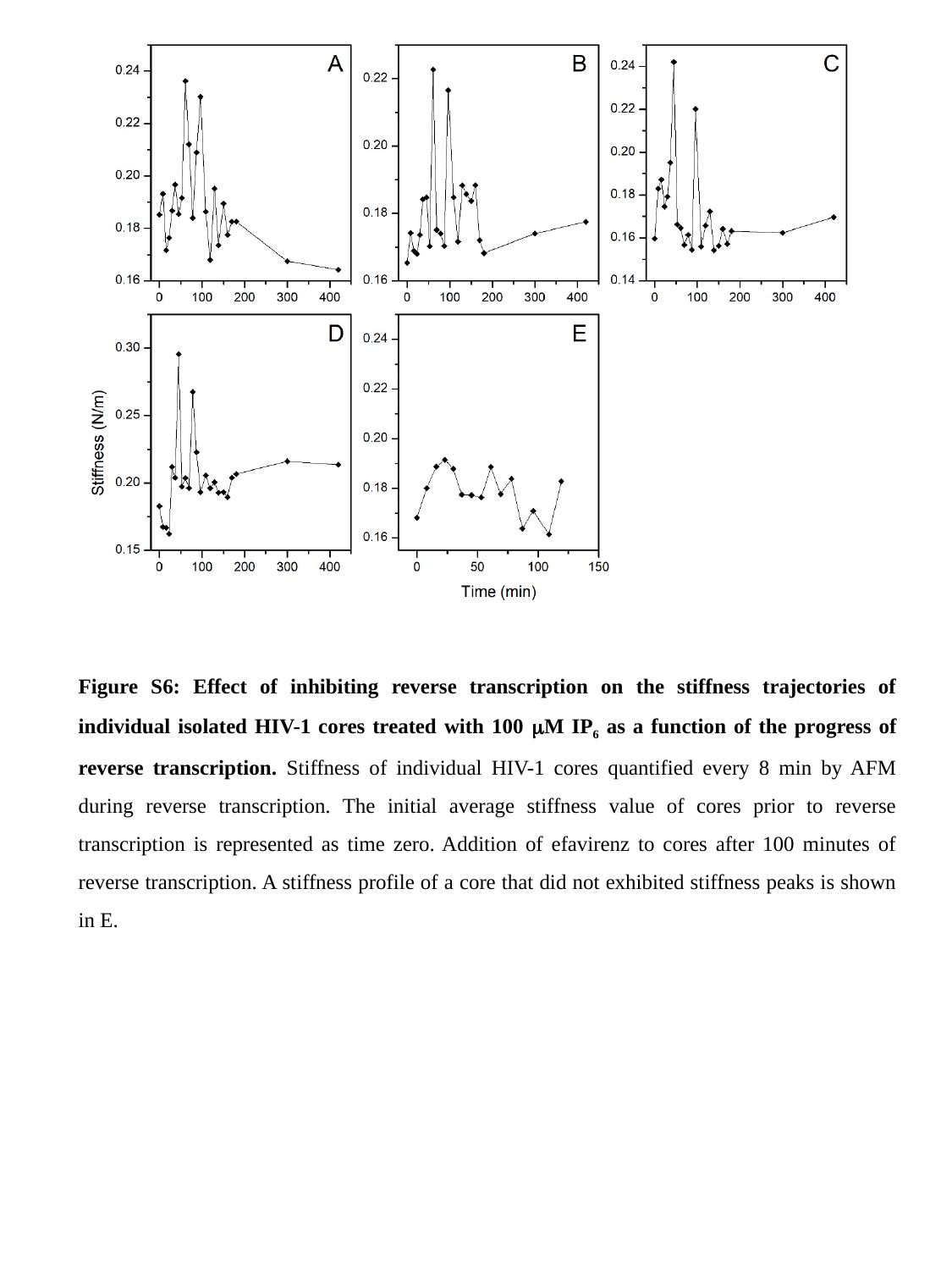

Figure S6: Effect of inhibiting reverse transcription on the stiffness trajectories of individual isolated HIV-1 cores treated with 100 μM IP_6_ as a function of the progress of reverse transcription. Stiffness of individual HIV-1 cores quantified every 8 min by AFM during reverse transcription. The initial average stiffness value of cores prior to reverse transcription is represented as time zero. Addition of efavirenz to cores after 100 minutes of reverse transcription. A stiffness profile of a core that did not exhibited stiffness peaks is shown in E.

### Slide 7
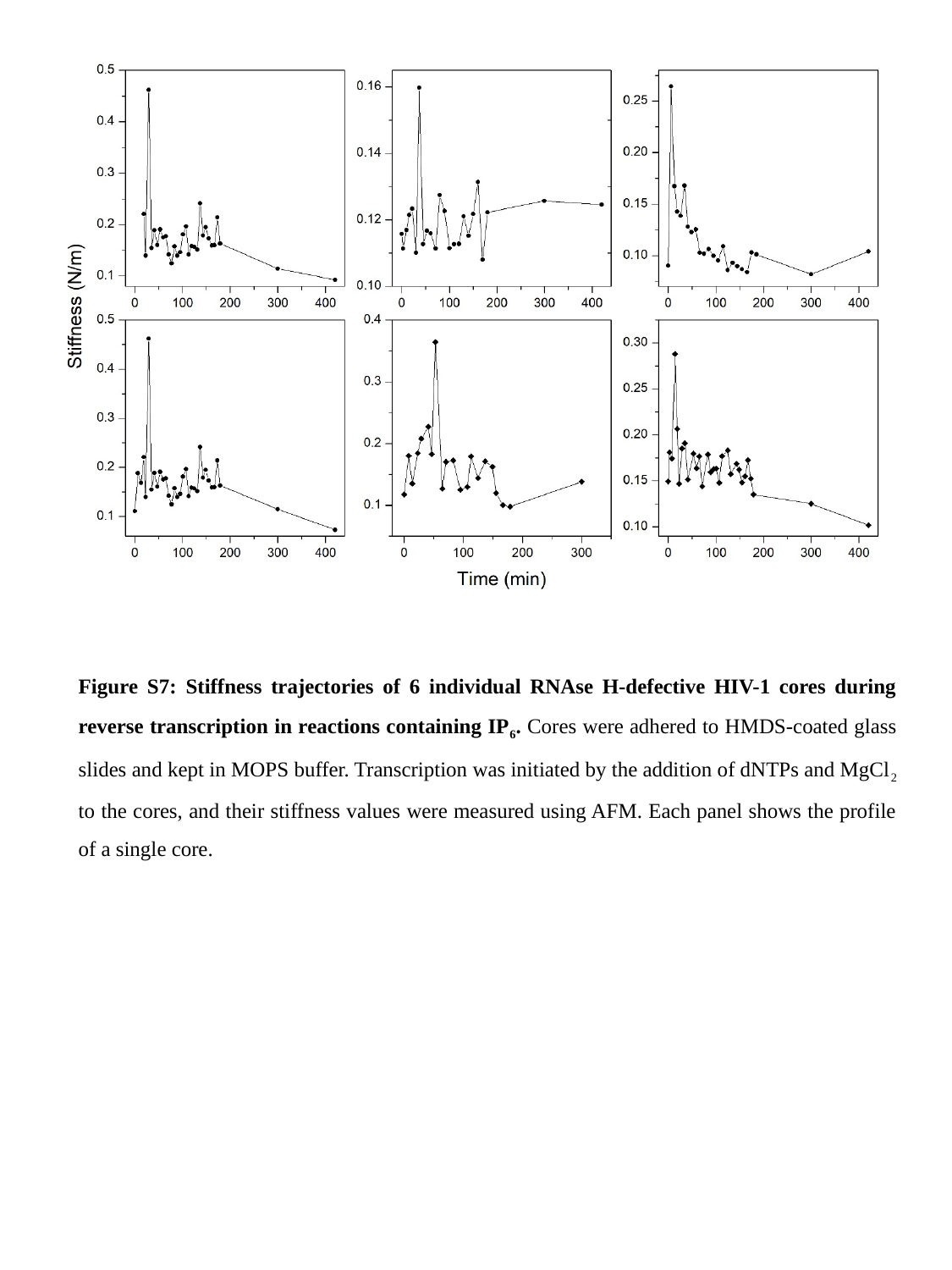

Figure S7: Stiffness trajectories of 6 individual RNAse H-defective HIV-1 cores during reverse transcription in reactions containing IP_6_. Cores were adhered to HMDS-coated glass slides and kept in MOPS buffer. Transcription was initiated by the addition of dNTPs and MgCl_2_ to the cores, and their stiffness values were measured using AFM. Each panel shows the profile of a single core.

### Slide 8
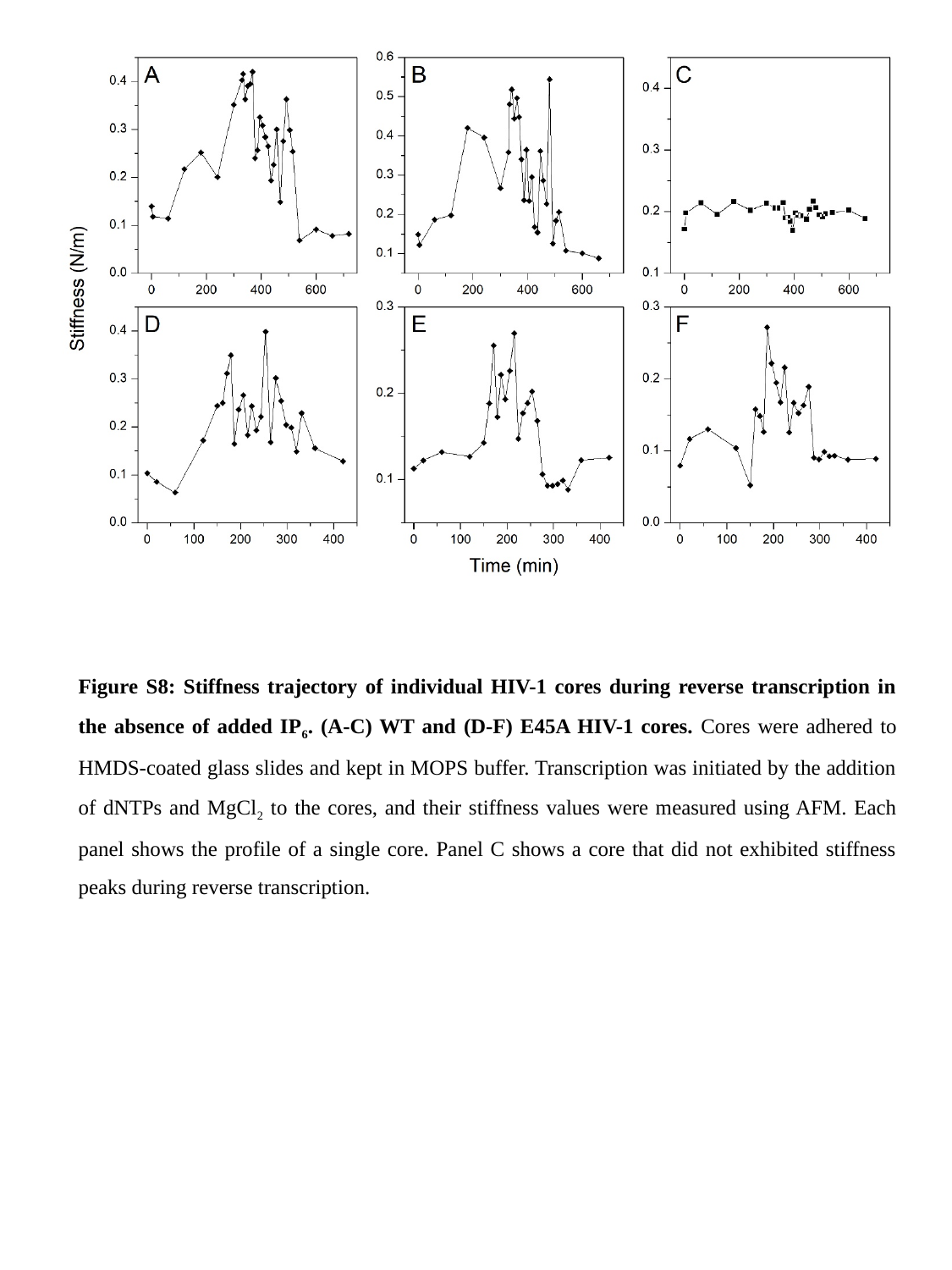

Figure S8: Stiffness trajectory of individual HIV-1 cores during reverse transcription in the absence of added IP_6_. (A-C) WT and (D-F) E45A HIV-1 cores. Cores were adhered to HMDS-coated glass slides and kept in MOPS buffer. Transcription was initiated by the addition of dNTPs and MgCl_2_ to the cores, and their stiffness values were measured using AFM. Each panel shows the profile of a single core. Panel C shows a core that did not exhibited stiffness peaks during reverse transcription.

### Slide 9
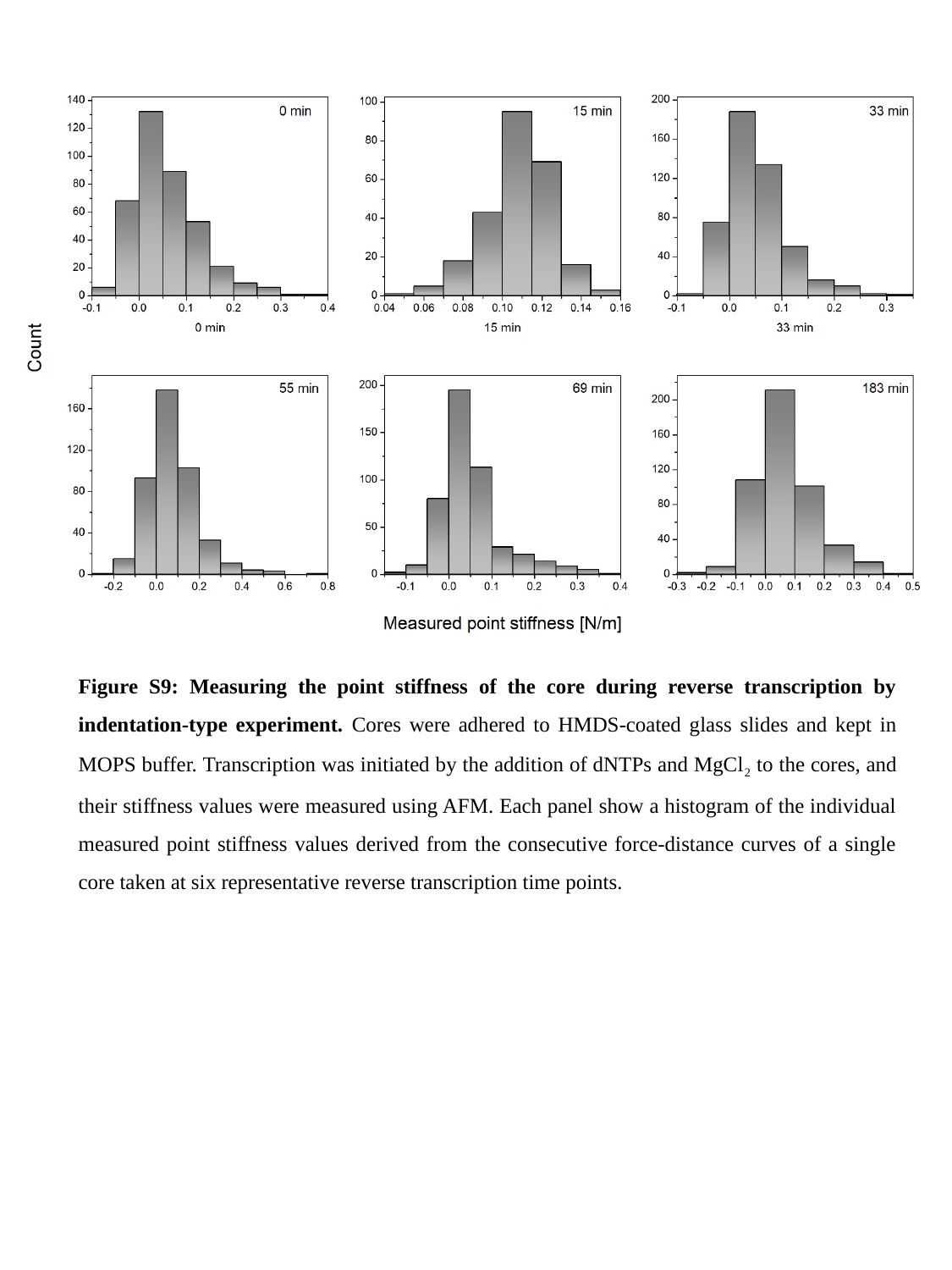

Figure S9: Measuring the point stiffness of the core during reverse transcription by indentation-type experiment. Cores were adhered to HMDS-coated glass slides and kept in MOPS buffer. Transcription was initiated by the addition of dNTPs and MgCl_2_ to the cores, and their stiffness values were measured using AFM. Each panel show a histogram of the individual measured point stiffness values derived from the consecutive force-distance curves of a single core taken at six representative reverse transcription time points.
